# Deconvolution of Cell Type-Specific Drug Responses in Human Tumor Tissue with Single-Cell RNA-seq

**DOI:** 10.1101/2020.04.22.056341

**Authors:** Wenting Zhao, Athanassios Dovas, Eleonora Francesca Spinazzi, Hanna Mendes Levitin, Pavan Upadhyayula, Tejaswi Sudhakar, Tamara Marie, Marc L. Otten, Michael Sisti, Jeffrey N. Bruce, Peter Canoll, Peter A. Sims

## Abstract

Precision oncology requires the timely selection of effective drugs for individual patients. An ideal platform would enable rapid screening of cell type-specific drug sensitivities directly in patient tumor tissue and reveal strategies to overcome intratumoral heterogeneity. Here we combine multiplexed drug perturbation in acute slice culture from freshly resected tumors with single-cell RNA sequencing (scRNA-seq) to profile transcriptome-wide drug responses. We applied this approach to glioblastoma (GBM) and demonstrated that acute slice cultures from individual patients recapitulate the cellular and molecular features of the originating tumor tissue. Detailed investigation of etoposide, a topoisomerase poison, and the histone deacetylase (HDAC) inhibitor panobinostat in acute slice cultures revealed cell type-specific responses across multiple patients, including unexpected effects on the immune microenvironment. We anticipate that this approach will facilitate rapid, personalized drug screening to identify effective therapies for solid tumors.

---

Inter- and intra-tumoral heterogeneity present major challenges for cancer therapy. Precision medicine, or targeted therapy, entails the use of agents that preferentially target tumor cells based on unique molecular features. The success of this approach relies on extensive characterization of tumor heterogeneity and microenvironment. While scRNA-seq can determine the cellular composition of complex tumors and even reveal cell type-specific drug sensitivities, these measurements are ultimately limited by models of drug response. Acute slice cultures are an attractive approach to modeling drug response in solid tumors because multiple cultures can be rapidly generated from a single surgical specimen, and they do not require extensive culturing or manipulation, which leads to distortion of the native composition of the tissue, selection, and loss of heterogeneity by diluting populations that do not proliferate rapidly^1-3^. Furthermore, drug perturbation experiments in acute slice cultures can be carried out rapidly, beginning on the day of surgical resection, on timescales relevant for clinical decision-making. GBM is an ideal setting for testing this approach because it exhibits profound inter- and intratumoral heterogeneity and is the most common and deadly primary brain malignancy in adults. Surgical resection is part of the standard-of-care, and robust protocols for acute slice culture of human GBM have been established previously^4-6^. Furthermore, GBM has been extensively characterized by scRNA-seq providing a detailed baseline for both the transformed populations that co-occur in individual patients and the microenvironment^7-11^. Indeed, single-cell characterization of GBM models has highlighted the importance of the tumor microenvironment in maintaining the phenotypic diversity of malignant cells^12^. We obtained GBM surgical specimens from six patients, generated multiple 500 micron slices, and placed them in short-term culture for drug perturbation (**Fig. 1A**). Screens were completed within 24 hours of surgery and analyzed immediately by scRNA-seq using our scalable microwell platform^13,14^ to deconvolve cell type-specific responses to multiple drugs (**Fig. 1A**).

**Figure 1.**
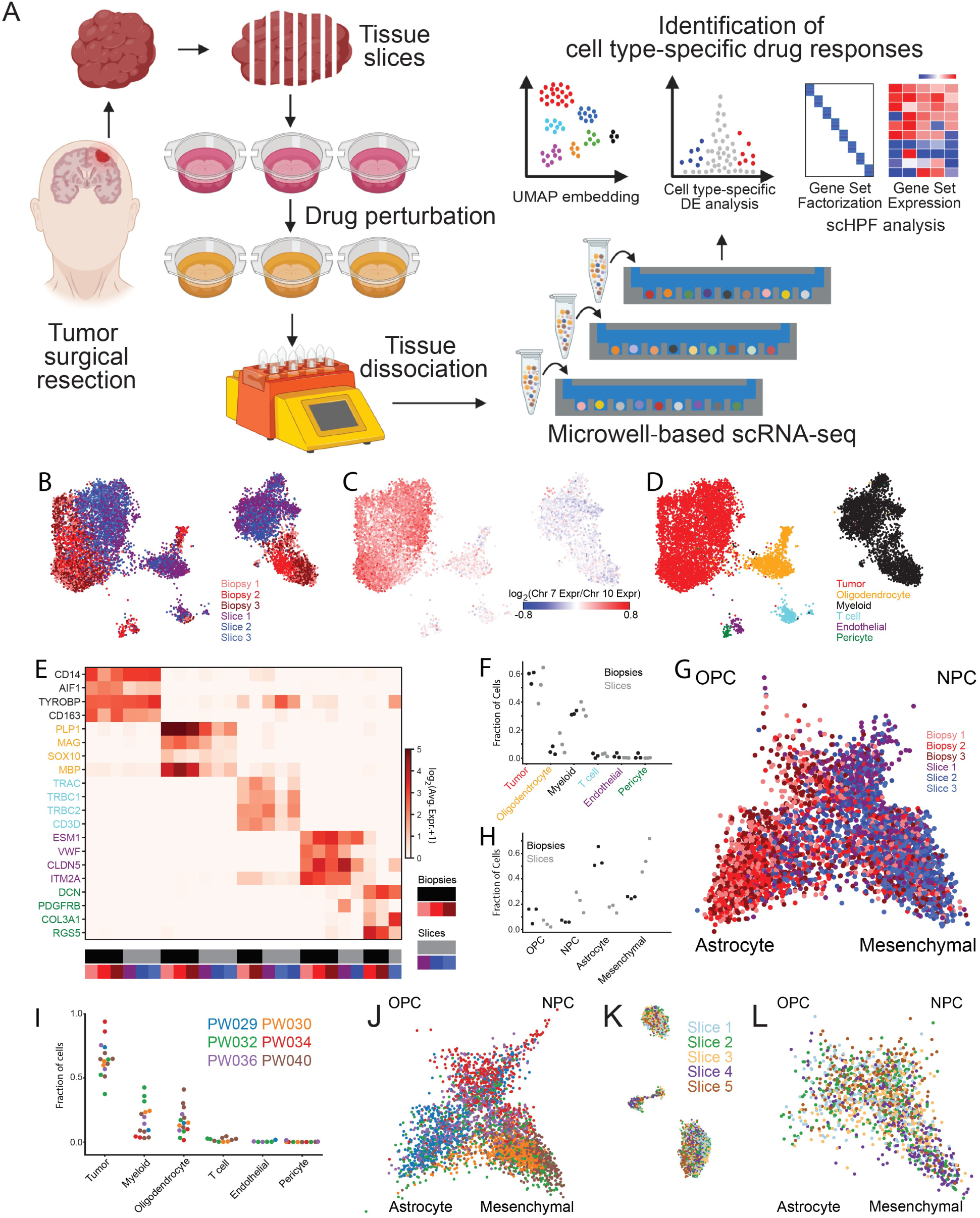
A) Schematic illustration of experimental and analytical methods for slice culture drug perturbation and scRNA-seq. B) UMAP embedding of scRNA-seq profiles from acutely isolated biopsies and slice cultures from different regions of the same tumor (PW032) colored by sample origin. C) Same as B) but colored by the log-ratio of Chr. 7 to Chr. 10 average expression where a high ratio (red) indicates malignant transformation. D) Same as B) but colored by cell type. E) Heatmap of average expression of marker genes from cell types in the tumor microenvironment in each cell type and sample from PW032. F) Fractional abundance of each major cell type in each biopsy and slice culture sample from PW032. G) Two-dimensional model projecting each transformed cell from PW032 biopsies and slice into four major GBM transformed populations colored by sample origin. I) Same as F) but for all untreated slice culture scRNA-seq data sets from the six patients in the study. J) Same as G) but for the transformed cells in all untreated slice culture scRNA-seq data sets from the six patients in the study. K) UMAP embedding of scRNA-seq profiles from five untreated slice cultures taken within 3.5 mm of each other from PW040 colored by sample of origin. L) Same as G) but for the transformed cells from the five untreated slice cultures from PW040.

We first demonstrated that acute slice culture preserves the cellular heterogeneity of GBM using scRNA-seq data from three uncultured biopsy specimens and three cultured slices obtained from the same patient (PW032). To identify subpopulations, we performed unsupervised clustering as previously reported^15,16^ on the entire data set containing 10,480 cells (4,358 from uncultured biopsies; 6,122 from cultured slices) and embedded the profiles in two dimensions using Uniform Manifold Approximation and Projection (UMAP, **Fig. 1B-D**)^17^. Transformed and untransformed subpopulations or clusters were distinguished by chromosome 7 amplification and chromosome 10 deletion, which were supported by both the scRNA-seq and whole genome sequencing (WGS) data (**Figs. 1C, Fig. S1A**, see Methods). By identifying highly enriched marker genes for each cluster, the non-malignant cells were further classified into myeloid cells (CD14, AIF1, TYROBP, CD163), oligodendrocytes (PLP1, MBP, MAG, SOX10), T cells (TRAC,

TRBC1, TRBC2, CD3D), endothelial cells (ESM1, ITM2A, VWF, CLDN5), and pericytes (PDGFRB, DCN,COL3A1, RGS5) (**Fig. 1D-E**). We observed transformed cells and all untransformed cell types with similar fractional composition (**Fig. 1F**) along with expression of their marker genes (**Fig. 1E**) in both uncultured biopsy samples and the cultured slices. Although the biopsies and slice cultures are similar, they are not identical, and systematic shifts in gene expression are evident in the embedding in **Fig. 1B**. Cell type-specific analysis in **Fig. S1B-D** highlights the major gene expression differences between the biopsies and slice cultures which could result from both spatial heterogeneity and culture conditions.

In previous studies, we used scRNA-seq to show that transformed cells in high-grade gliomas resemble oligodendrocyte lineage cells (including progenitors or OPCs), astrocytes, neuronal precursors, and mesenchymal cells at the level of gene expression^9^, consistent with earlier work using bulk analysis^18-20^. More recently, Neftel *et al.* developed an elegant model based on gene signatures of these four major states to classify scRNA-seq profiles of glioma cells^11^. We used this model to examine the transformed cells in the biopsies and slice cultures in detail and found that all four states were well-represented in both the biopsy and slice cultures, and that most cells classified as astrocyte-like or mesenchymal in this particular tumor (**Fig. 1G-H**). However, the slice cultures contained more mesenchymal cells whereas the biopsy cells were more astrocyte-like (**Fig. 1G-H**). While this could be due, in part, to culture conditions, we expect this level of variation based on previous studies of spatial heterogeneity since the slice cultures and biopsies were obtained from different regions of the tumor^8,15,19,21^. We conducted a similar analysis of slice cultures from six patients (including PW032) and found representation of transformed cells (**Fig. S2**) and the same major cell types with similar relative abundances after 24 hours of culture (**Fig. 1I**). We also analyzed their transformed populations using the Neftel *et al.* model (**Fig. 1J**)^11^, and found good representation of all four major GBM states with some tumors appearing more proneural (OPC/NPC – PW034) and others more astrocytic (PW029), mesenchymal (PW030, PW040), or both (PW032, PW036) as quantified in **Fig. S2D**. Finally, to analyze spatial effects across slice cultures within a single resection, we profiled five slice cultures such that each was 500 μm thick and the interval between the two most adjacent slices was also 500 μm (maximum spatial distance of 3.5 mm). scRNA-seq profiles of the five slices co-clustered well based on UMAP embedding (**Fig. 1K, Fig. S3**) and the four-state model of the transformed cells (**Fig. 1L**) and showed good representation of major cell types (**Fig. S3**). Taken together, these data suggest that cultured slices preserve the major cellular and molecular features of the tumor microenvironment and represent the well-established inter- and intra-tumoral heterogeneity observed in gliomas.

To test the feasibility of personalized drug screening with patient-derived slice cultures and scRNA-seq, we perturbed slices derived from one GBM resection (PW030) with six different drugs chosen for diverse mechanisms of action and included two vehicle controls (**Fig. 2A** and **Table S1**). We profiled 48,404 cells from eight slices and identified transformed and untransformed populations as described above. To identify drug-induced transcriptional changes, we performed differential expression analysis for the tumor, myeloid and oligodendrocyte populations (**Fig. 2B-C**). Treatment with histone deacetylase (HDAC) inhibitor panobinostat resulted in the strongest response with 9,632, 4,228, and 3,183 significantly differentially expressed genes (p<0.01) in the tumor, myeloid, and oligodendrocyte populations (**Fig. 2B**), respectively, with similar results when we restricted our analysis to fold-changes >2 (**Fig. 2C**). To identify drugs with highly specific effects on subpopulations of tumor cells, we first computed a UMAP embedding for the transformed cells from the control slices (**Fig. 2D**), which we use as a reference for comparison to treated cells. The majority of control tumor cells appear mesenchymal (**Fig. 1J**) with pervasive expression of CD44 and VIM and an astrocytic subpopulation expressing GFAP at high levels (**Fig. 2D**). However, there is a small subpopulation of proliferating cells marked by TOP2A and MKI67 (**Fig. 2D**). Next, we projected the profiles of transformed cells from each treated slice into this embedding (**Fig. 2E**). Consistent with the differential expression analysis, we observed that panobinostat had a dramatic compositional effect on the transformed cells. We also noticed that etoposide selectively eliminated the small, proliferative subpopulation, consistent with its mechanism-of-action as a topoisomerase poison^22^. Given the disparate effects of these two drugs in PW030, we made them the focus of our subsequent analysis.

**Figure 2.**
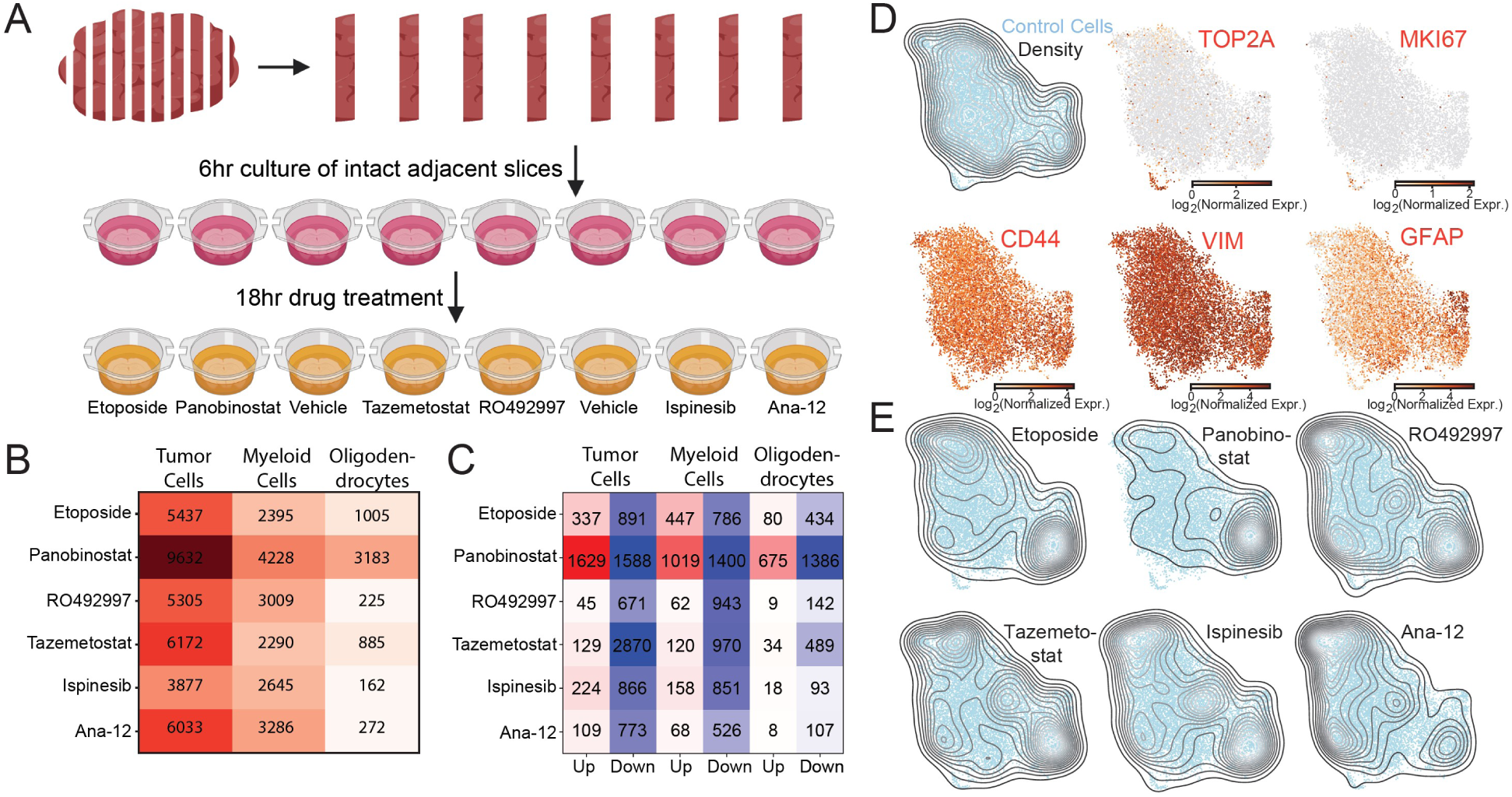
A) Experimental schematic for slice culture drug screening (6 drugs, 2 controls) from a single patient (PW030). B) Heatmap showing the number of differentially expressed genes (FDR<0.01) in the tumor, myeloid, and oligodendrocyte populations between treated and control slices for each drug in the screen illustrated in A). C) Same as B) but showing only differentially expressed genes with FDR<0.01 and fold-change amplitude greater than two (both up- and down-regulated genes). D) UMAP embedding of scRNA-seq profiles of transformed cells from the control slices colored by expression of two proliferation markers (TOP2A, MKI67), two mesenchymal markers (CD44, VIM), and an astrocyte marker (GFAP). E) Same as D) but with UMAP projection density of scRNA-seq profiles of transformed cell from the treated slice cultures for each drug. Note that there is negligible projection density for the etoposide-treated cells onto the control cells for the small proliferative population expressing TOP2A and MKI67.

To identify cell type-specific responses to etoposide and panobinostat that are conserved across patients, we conducted slice culture drug-perturbations across six GBM patients followed by scRNA-seq. **Table S2** contains a summary of all the slice culture samples used for this analysis. After subsampling the scRNA-seq profiles from each vehicle- and drug-treated slice culture from all six patients to the same number of cells, we generated a low-dimensional representation of the merged data using single-cell hierarchical Poisson factorization (scHPF)^15^. This Bayesian algorithm operates directly on the count matrix and identifies latent factors corresponding to the major gene expression programs that define the population. We identified 15 factors associated with canonical markers of neural cell types, GBM subpopulations, biological processes (e.g. proliferation), and drug response (**Fig. S4, Table S4**). Two nuisance factors were associated with coverage (enriched in ribosomal and other housekeeping genes) and cell stress (e.g. heat shock – likely a dissociation artifact) and removed from the model (**Fig. S4A, Table S4**).

To visualize the model, we created a UMAP embedding of the scHPF cell score matrix (**Fig. 3A-F**, see Methods). Based on aneuploidy analysis of chromosomes 7 and 10, the transformed cells from each patient separate into essentially non-overlapping clusters (**Fig. 3A,C**). In contrast, untransformed oligodendrocytes, myeloid cells, and T cells overlap significantly across the six patients (**Fig. 3A,D-F**), consistent with previous studies of fresh resections^9^. We also observed that panobinostat- and vehicle-treated cells generally showed little overlap across all cell types, while etoposide-treated cells tended to co-cluster with the controls (**Fig. 3B**). This is consistent with our screening results above which suggest that panobinostat significantly alters gene expression, whereas etoposide primarily impacts genes involved in proliferation.

**Figure 3.**
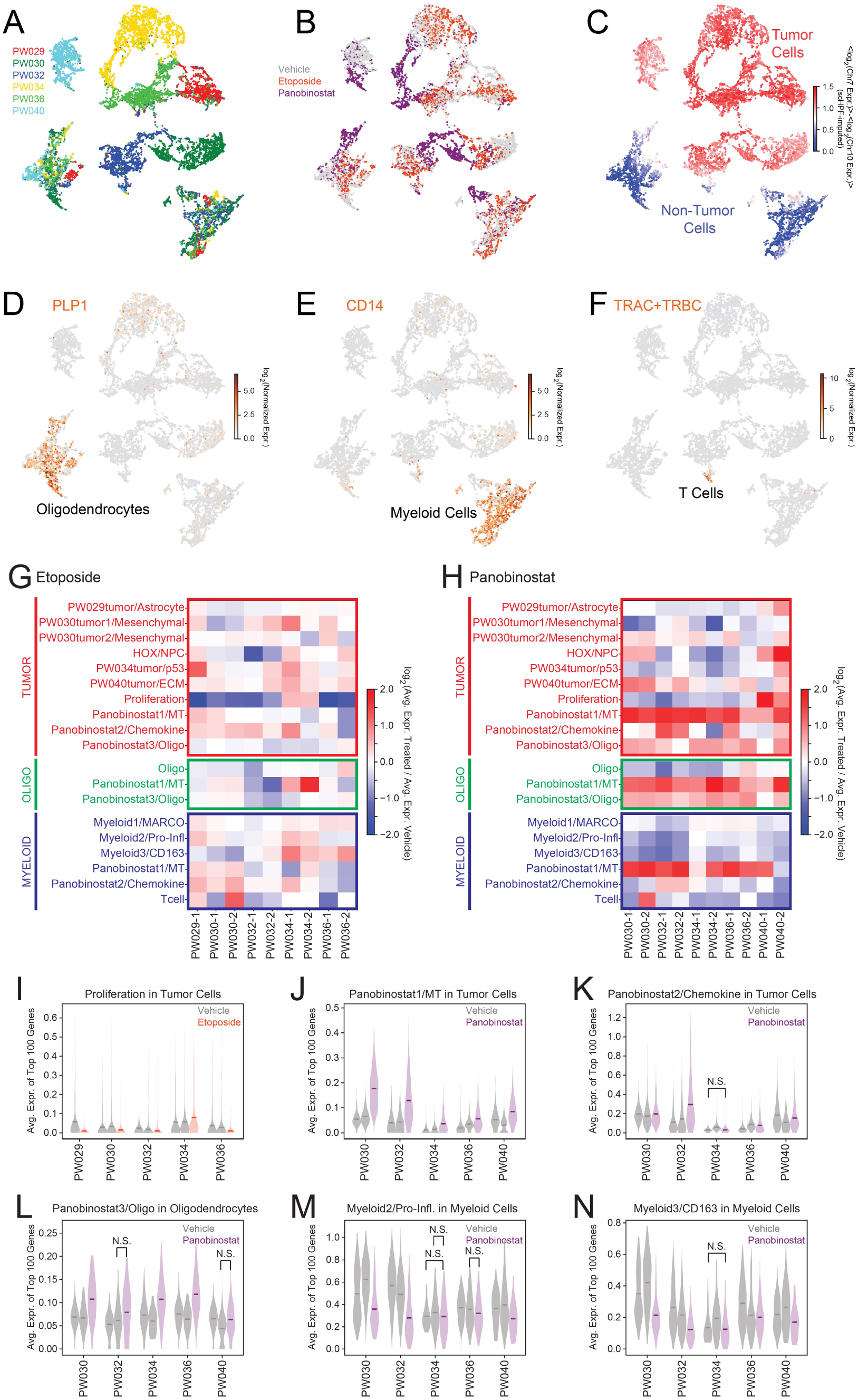
A) UMAP embedding of scRNA-seq profiles from slice cultures of six patients generated using the cell score matrix from joint scHPF analysis of the entire data set colored by patient. B) Same as A) but colored by treatment condition. C) Same as A) but colored by the scHPF-imputed log-ratio of Chr. 7 to Chr. 10 average expression where a high ratio (red) indicates malignant transformation. D) Same as A) but colored by expression of the oligodendrocyte marker PLP1. E) Same as A) but colored by expression of the myeloid marker CD14. F) Same as A) but colored by the total expression of the T cell receptor constant regions (TRAC, TRBC1, TRBC2). G) Heatmap showing the log-ratio of the average expression of the top 100 genes in each eptoposide-treated to each control slice for each scHPF factor and each of three cell types – transformed (tumor), oligodendrocyte (oligo), and myeloid. H) Same as G) for panobinostat-treated slices. I) Violin plots showing the distributions of the average expression of the top 100 genes in the Proliferation scHPF factor for each vehicle- and etoposide-treated slice for each patient in tumor cells. All within-patient, vehicle-treatment comparisons have p<0.05 (Mann-Whitney U-test) unless otherwise indicated (N.S. or not significant). J) Same as I) for the Panobinostat1/MT scHPF factor for each vehicle- and panobinostat-treated slice. In tumor cells K) Same as J) for the Panobinostat2/Chemokine scHPF factor in tumor cells. L) Same as J) for the Panobinostat3/Oligo scHPF factor in oligodendrocytes. M) Same as J) for the Myeloid2/Pro-Inflammatory scHPF factor in myeloid cells. N) Same as J) for the Myeloid3/CD163 scHPF factor in myeloid cells.

To identify conserved drug responses, we compared the expression of top genes in each factor between the drug- and vehicle-treated slices from each patient. As expected, the most conserved response to etoposide was a decrease in expression of the Proliferation factor in the tumor compartment (**Fig. 3G,I**). This occurred in all but one patient, PW034, despite its high levels of cycling cells (**Fig. 3I**). Etoposide did not show consistent effects on other factors and had limited impact overall on oligodendrocytes and myeloid cells (**Fig. 3G**). We validated the loss of TOP2A^+^ tumor cells, which we also observed by conventional differential expression analysis (**Fig. S5A**), using *in situ* hybridization analysis of etoposide-treated slice cultures from a separate cohort (**Fig. S6A**). In contrast, panobinostat affected multiple factors for both tumor and non-tumor cells (**Fig. 3H**). We observed a modest decrease in expression for the Proliferation factor across all patients except for PW040 (**Fig. 3H**). Interestingly, panobinostat induced expression of LEFTY1, BEX5, and SAXO2 as part of the Panobinostat3/Oligo factor, which was predominantly oligodendrocyte-specific (**Fig. 3H, Fig. S4B**). However, the most notable effect was upregulation of metallothionein family genes (Panobinostat1/MT factor) across all cell types (**Fig. 3H,J**), consistent with previous reports that HDAC inhibitors can perturb this highly inducible gene cluster^23,24^. Interestingly, cell type-specific differential expression analysis not only confirmed metallothionein induction but also revealed upregulation of mature neuronal genes (e.g. SNAP25, SLC17A7, KCNB1, RAB3A), a component of the same factor, specifically in tumor cells (**Fig. S5C**). Because we observed metallothionein induction in all six patients and the three major cell populations analyzed here, it is a potentially useful biomarker of panobinostat response.

Panobinostat treatment significantly impacted gene expression in myeloid cells. In slice cultures from 3/5 patients, we observed a modest decrease in a factor marked by pro-inflammatory cytokines (**Fig. 3H,M**), which have been shown to be predominantly expressed by microglia in the glioma microenvironment^9,25^. We observed a more consistent effect on a myeloid factor marked by CD163, which is likely expressed by macrophages which are thought to be immunosuppressive in GBM (**Fig. 3H,N**)^25^. However, a third myeloid factor marked by MARCO, which has been associated with mesenchymal transformation and poor survival in GBM (Chen *et al*, submitted manuscript enclosed)^26^, was relatively unaffected by panobinostat (**Fig. 3H**). We verified that CD163 exhibited significant, myeloid-specific differential expression (**Fig. S5D**) and the loss of CD163^+^ macrophages in general and relative to CCL3^+^ pro-inflammatory myeloid cells by *in situ* hybridization analysis of vehicle and panobinostat-treated slice cultures from a separate group of patients (**Fig. S6B**).

Collectively, this work establishes a multiplexed experimental and analytical pipeline to deconvolute cell type-specific drug response in GBM tissue from individual patients. Acute slices generated from fresh tumor tissues preserve the key molecular and cellular features of the original tissue, and provide a setting for drug response to be evaluated on multiple tumor cell subpopulations and cell types in the tumor microenvironment. We further demonstrated the feasibility of conducting personalized drug screens using this approach with a turnaround time of less than one week after surgery – a relevant timescale for clinical decision-making. Focused analysis of etoposide and panobinostat across six patients (five for each drug) revealed drug-induced responses in specific populations of transformed and microenvironmental cells, patient specific drug sensitivities, and drug effects conserved across patients. Etoposide consistently downregulated cell cycle genes in proliferating tumor cells with minimal conserved effects on untransformed or less proliferative transformed cells. The HDAC inhibitor panobinostat induced the expression of metallothionein family genes and mature neuronal genes in tumor cells, and significantly re-modeled the myeloid population in the tumor microenvironment. Overall, we hope that this approach will find broad utility for pre-clinical studies and potentially for rapid, personalized drug screening to develop cellular and molecular enrollment criteria for clinical trials.

## Methods

### Preparation and culture of tissue slices

This work was approved by the Columbia University Irving Medical Center Institutional Review Board before commencing the study. Patient diagnosis information can be found in **Table S2**. Tumor specimens were collected immediately after surgical removal and kept in ice-cold artificial cerebrospinal fluid (ACSF) solution containing 210 mM sucrose, 10 mM glucose, 2.5 mM KCl, 1.25 mM NaH_2_PO_4_, 0.5 mM CaCl_2_, 7 mM MgCl_2_ and 26 mM NaHCO_3_ for transportation. Preparation of *ex vivo* tissue slices was modified from methods described previously^5^. Briefly, the collected tumor specimens were placed in a drop of ice-cold ACSF and sliced using a tissue chopper (McIlwain) at a thickness of 500 µm under sterile conditions. The slices were immediately transferred to the ice-cold ACSF solution in 6-well plates using a sterile plastic Pasteur pipette half filled with ice-cold ACSF solution followed by a 15 minutes recovery in ACSF to reach room temperature. Intact and well-shaped slices (approximately 5-10 mm diameter) were then placed on top of a porous membrane insert (0.4 µm, Millipore). Then the membrane inserts were placed into 6-well plates containing 1.5 mL maintenance medium consisting of F12/DMEM (Gibco) supplemented with N-2 Supplement (Gibco) and 1% antibiotic-antimycotic (ThermoFisher). To ensure proper diffusion into the slice, culture medium was placed under the membrane insert without bubbles. A drop of 10 µl of culture medium was added directly on top of each slice to prevent the slice surface from drying. The slices were first rested for 6 hrs with the maintenance medium in a humidified incubator at 37°C and 5% CO_2_. Then, the medium was replaced with pre-warmed medium containing drugs with desired concentration (**Table S1**) or corresponding volume of vehicle (DMSO). Slices were then cultured with the treatment medium in a humidified incubator at 37°C and 5% CO_2_ for 18 hrs before being collected for dissociation.

### Dissociation of tissue and slices

Collected tissue samples or tissue slices were dissociated using the Adult Brain Dissociation kit (Miltenyi Biotec) on gentleMACS Octo Dissociator with Heaters (Miltenyi Biotec) according to the manufacturer’s instructions.

### Microwell scRNA-seq

Dissociated cells from each slice were profiled using microwell-based single-cell RNA-seq^14^ as previously described^9,15^ with the following modifications: once the RNA-capture step was finished, sealing oil was flushed out of the devices by pipetting 1 mL of wash buffer supplemented with 0.04 U/μl RNase inhibitor (Thermo Fisher Scientific) and then beads were extracted from the device and resuspended in 200 µl of reverse transcription mixture. Bead-suspensions were divided into 50 µl aliquots and placed into PCR tubes (Corning) followed by incubation at 25°C for 30 min and at 42°C for 90 min in a thermocycler. Each cDNA library was barcoded with an Illumina sample index. Libraries with unique Illumina sample indices were pooled for sequencing on 1) an Illumina NextSeq 500 with an 8-base index read, a 21-base read 1 containing cell-identifying barcodes (CB) and unique molecular identifiers (UMIs), and a 63-base read 2 containing the transcript sequence, or 2) an Illumina NovaSeq 6000 with an 8-base index read, a 26-base read 1 containing CB and UMI, and a 91-base read 2 containing the transcript sequence.

### scRNA-seq data preprocessing

Raw data obtained from the Illumina NextSeq 500 was trimmed and aligned as described previously^9^. For each read with a unique, strand-specific alignment to exonic sequence, we constructed an address comprised of the CB, UMI barcode, and gene identifier. Raw data obtained from the Illumina NovaSeq 6000 was first corrected for index swapping to avoid cross-talk between sample index sequences using the algorithm described by Griffiths *et al*^27^ before assigning read addresses for each sample. For samples had been sequenced on both Illumina NextSeq 500 and NovaSeq 6000, we combined the addresses from the NextSeq 500 and the corrected addresses from the NovaSeq 6000 for data processing as described previously^9,15^. Briefly, reads with the same CB, UMI and aligned gene were collapsed and sequencing errors in the CB and UMI were corrected to generate a preliminary matrix of molecular counts for each cell.

We applied the EmptyDrops algorithm to recover truecell-identifying barcodes in the digital gene expression matrix^28^. We then removed CBs that satisfied any of the following criteria: 1) fractional alignment to the mitochondrial genome greater than 1.96 standard deviations above the mean; 2) a ratio of molecules aligning to whole gene bodies (including introns) to molecules aligning exclusively to exons greater than 1.96 standard deviations above the mean; 3) average number of reads per molecule or average number of molecules per gene >2.5 standard deviations above the mean for a given sample; or 4) more than 40% of UMI bases are T or where the average number of T-bases per UMI is at least 4..

### Unsupervised clustering, differential expression, and visualization

Clustering, visualization, and identification of cluster-specific genes was performed as described previously (www.github.com/simslab/cluster_diffex2018)^15^. We used Louvain community detection as implemented in Phenograph for unsupervised clustering with k=20 for all k-nearest neighbor graphs^16^. For all clustering and visualization analyses of merged datasets, we first identified marker genes using the drop-out curve method described in Levitin *et al*.^15^ (www.github.com/simslab/cluster_diffex2018) for each individual sample and took the union of the resulting marker sets to cluster and embed the merged dataset. We projected drug-treated cells onto vehicle-treated cells with UMAP in **Fig. 2** as described in Szabo *et al* (code available at www.github.com/simslab/umap_projection)^29^.

### Whole genome sequencing and analysis

Genomic DNA was extracted from a piece of frozen tissue from each tumor using the DNeasy Blood & Tissue Kits (Qiagen) according to the manufacturer’s instructions and was submitted to the Beijing Genomics Institute (BGI) for whole genome sequencing using their DNBseq technology. Raw sequencing data were aligned to the human genome using bwa mem and analyzed as described in Yuan *et al*^9^. Briefly, we computed the number of de-duplicated reads that aligned to each chromosome for each patient and divided this by the number of de-duplicated reads that aligned to each chromosome for a diploid germline sample from patient PW034 (pooled blood mononuclear cells) after normalizing both by total reads. We then normalized this ratio by the median ratio across all chromosomes and multiplied by two to estimate the average copy number of each chromosome.

### Identification of malignant glioma cells and non-tumor cells

Chr. 7 amplification and Chr. 10 deletion were observed from the whole-genome sequencing results for all patients in this cohort. Therefore, we identified the transformed cells and untransformed cells using a linear combination of normalized average chromosome 7 and 10 expression in each cell as follows. We first merged scRNA-seq data of all samples derived from the same patient for unsupervised clustering analysis and defined putative malignant cells and non-tumor cells using the genes most specific to each cluster. Putative tumor-myeloid doublet clusters were removed prior to malignant analysis. Next, we computed the average gene expression on each somatic chromosome as described in Yuan *et al*^9^. We define the malignancy score to be the log-ratio of the average expression of Chr. 7 genes to that of Chr. 10 genes and plotted the distribution of malignancy score. We fit a double Gaussian to the malignancy score distribution and established a threshold at 1.96 standard deviations below the mean of the Gaussian with the higher mean (i.e. 95% confidence interval). Putative malignant cells with malignancy scores below this threshold and putative non-tumor cells with malignancy scores above this threshold were discarded as non-malignant or potential multiplets.

For the comparison of biopsy and acute slice cultures from PW032 shown in **Fig. 1**, we co-clustered all of the samples together using Phenograph and identified a cluster that was statistically enriched in genes associated with red blood cells (HBA1, HBA2, HBB), transformd glioma cells (SAA1, GFAP), and myeloid cells (CD14, C1QA). We discarded these cells as potential multiplets before completing our analysis. Similarly, for the drug screening analysis of PW030 in **Fig. 2**, we co-clustered all eight samples using Phenograph and identified a cluster that was statistically enriched in markers of transformed glioma cells (GFAP) and myeloid cells (CD14). We discarded these cells as potential multiplets before completing our analysis.

### Single cell hierarchical Poisson factorization (scHPF) analysis

For the scHPF model in Fig. 3, we combined scRNA-seq profiles from one vehicle-treated and one etoposide-treated slice from PW029; two vehicle-treated, one etoposide-treated, and one Panobinostat-treated slice from PW030, PW032, PW034, and PW036; and two vehicle-treated and one Panobinostat-treated slice from PW040 for a total of 21 samples (see **Table S2**). To avoid dominant factors from any one sample, we randomly sub-sampled the scRNA-seq profiles such that each of the 21 samples contributed 803 cells to the model for a total of 16,863 cells. We then factorized the resulting merged count matrix using scHPF with default parameters and K = 17 (www.github.com/simslab/schpf)^15^. For all downstream analysis of the model, we removed two nuisance factors. The first was correlated with coverage and highly ranked housekeeping genes and ribosomal protein-encoding genes, and the second contained highly ranked genes associated with cell stress and heat shock, likely a result of dissociation artifacts in a subset of cells and samples (**Fig. S4A**). This resulted in a scHPF model with 15 factors (**Table S4**).

To visualize the scHPF model, we generated a UMAP embedding using a Pearson correlation matrix computed from the cell score matrix. To cluster the scRNA-seq profiles using the Phenograph implementation of Louvain community detection^16^, we used the same Pearson correlation matrix and k=50 to construct a k-nearest neighbors graph. We conducted the aneuploidy analysis in **Fig. 3C** from the scHPF model by first computing the cell loading matrix Θ containing elements *E*[*θ*_*i,k*_|*x*] for each cell-factor pair *i,k* and the gene sample weight matrix B containing elements *E*[*β*_*g,k*_|*x*] for each gene-factor pair *g,k* where *x* is the scRNA-seq count matrix. Next, we computed the diagonal cell scaling matrix Ξ containing elements *E*[*ξ*_*i,i*_|*x*]*10000 for each cell *i* and finally:

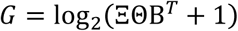

where *G* is the log-transformed scHPF-imputed expectation value matrix for the expression level of each gene in each cell. We colored the UMAP embedding in **Fig. 3C** by the difference in the average value of *G* for genes in chromosome 7 and that for chromosome 10. We scored each Phenograph cluster by the average of this value and took all cells in clusters with an above-average score to be malignantly transformed.

The fold-change values in the heatmaps in **Figs. 3G-H** were computed by dividing the average expression of the top 100 genes in each factor (rows) for the treated slice by that of each vehicle-treated slice (columns) and log-transforming. For select factors, the distribution of average expression of the top 100 genes across cells is shown for the tumor cells, oligodendrocytes, or myeloid cells for each slice in **Figs. 3I-N**.

### Cell-type specific differential expression analysis

To maximize our statistical power for the cell type-specific differential expression analysis shown in **Fig. S5**, we added back all of the scRNA-seq profiles that we had subsampled out of the data set when we constructed the scHPF model, as described above. To project these additional data onto our existing scHPF model, we held the variational distributions for global, gene-specific variables fixed while updating the variational distributions for cell-specific local variables as when the model was originally trained. We used the “prep-like” command in scHPF to select the same genes that were used in the original model, and then projected the data with the “project” command in scHPF with the ‘—recalc-bp’ option and default parameters. This results in variational approximations for the cell budgets *ξ*_*i*_ and weights *θ*_*i,k*_ for the additional, projected data, but does not alter the gene budgets *η*_*g*_ or weights *β*_*g,k*_, nor does it alter the cell budgets or weights for the cells used to train the original model. To associate the projected cells with the originally defined Phenograph clusters, we used the “classify” command in Phenograph^16^ with a Pearson correlation matrix derived from the cell score matrix computed by scHPF projection. This allowed us to assign the additional cells as transformed, myeloid, etc.

To identify differentially expressed genes for drug- vs. vehicle-treated tumor and myeloid cells, we first randomly sub-sampled the condition with a greater number of cells in each comparison to have the same number of cells as the condition with fewer cells. Next, we subsampled the count matrices for the two conditions such that they had the same average number of molecules per cell and normalized the resulting count matrix using scran^30^. We then conducted differential expression analysis for protein-coding genes using the two-sided Mann-Whitney U-test as implemented by the “mannwhitneyu” command in the Python module “scipy”. The resulting p-values were corrected for false discovery using the Benjamini-Hochberg procedure as implemented in the “mutlipletests” command in the Python module “statsmodels”. We note that this same approach was used for the differential expression analysis shown in **Fig. 2B-C**.

### Data and code availability

All sequencing data have been deposited in GEO XX. Processed data and basic association analyses will be made available upon request. The computer code for unsupervised clustering and visualization is available at https://github.com/simslab/cluster_diffex2018, scHPF is available at https://github.com/simslab/scHPF, and the UMAP projection code is available at https://github.com/simslab/umap_projection. All of the raw sequencing data and processed count matrices are available on the Gene Expression Omnibus under accession GSE148842.

### In situ hybridization and microscopy

To detect changes *in situ* of TOP2A, SOX2, CD163 and CCL3 mRNA upon etoposide and panobinostat treatment, we performed RNAscope on vehicle- or drug-treated slices from three separate cohorts that were not processed for scRNA-seq. Treated slices were fixed in 4% PFA overnight at 4°C, paraffin-embedded, and cut into 5 mm sections. Probes against the above-mentioned mRNAs were obtained from ACDBio (**Table S3**). *In situ* hybridization (ISH) was performed according to the manufacturer’s protocol for the RNAScope® Multiplex Fluorescent V2 Assay (ACDBio). Briefly, serial sections were baked at 60 °C for 1 hour before being deparaffinised in xylene and 100% ethanol. After drying the slides for 5 min at 60 °C, H_2_O_2_ was added for 10 min at RT. For antigen accessibility, slides were incubated in boiling 1X Target Retrieval reagents (∼98 °C) for 15 min, washed in water, dehydrated in 100% ethanol and finally treated with Protease Plus for 30 min at 40 °C. The C3 probes were diluted in C1 probes at a 1:50 ratio and incubated on the slides for 2 hrs at 40 °C. C1 probes were detected with TSA-Cy3 (Perkin Elmer, NEL744001KT) and C3 probes were detected with TSA-Cy5 (Perkin Elmer, NEL745001KT). DAPI was added to label the nuclei, and slides were mounted using Fluoromount. After drying at RT, the mounted slices were stored in the dark at 4 °C.

Images were acquired on a Zeiss LSM 800 confocal microscope with a 40×/1.3 NA oil immersion objective, using 405 nm, 561 nm and 639 nm excitation. Five to six fields per probe were selected based on high SOX2 expression in serial sections, a pervasive marker of transformed glioma cells^9^. Confocal stacks were acquired with a 1 Airy pinhole and at 0.58 μm steps. Images were exported to FIJI/ImageJ for further analysis. Maximum intensity projections were generated and each image was auto-thresholded using the method by Yen (TOP2A, SOX2 and CCL3) or Otsu (CD163) in order to eliminate background autofluorescence. Particles were counted and the integrated intensity value was extracted. TOP2A was expressed as integrated density over total number of cells per sample in control versus etoposide-treated sections. Changes in CD163 message were expressed as a ratio of CD163 integrated density over CCL3 integrated density in control versus panobinostat-treated samples.

## Competing Interests Statement

P.A.S. is listed as an inventor on patent applications filed by Columbia University related to the microwell technology described here.

## Contributions

W.Z. designed and executed the drug treatment and scRNA-seq experiments with input from P.A.S. and P.C. E.F.S., T.S., T.M., M.L.O., M.S., J.N.B., and P.C. procured the tissue. E.F.S. and P.U. generated tissue slices. A.D. and E.F.S. optimized the slice culture protocol with input from P.C. W.Z. and A.D. performed and analyzed the RNAscope experiments and imaging data. H.M.L. developed the scHPF projection analysis. W.Z. and P.A.S. performed computational analysis. W.Z. and P.A.S. wrote the manuscript. All authors edited, read and approved the final manuscript.

## Acknowledgements

We thank Erin C. Bush and Michael Finlayson in the Columbia Single Cell Analysis Core and Sulzberger Columbia Genome Center for technical support. P.A.S. was supported by the Mark Foundation for Cancer Research Grant MFCR18-019 ELA. J.N.B, P.C., and P.A.S. were supported by NIH/NINDS R01NS103473. This research was funded in part through the NIH/NCI Cancer Center Support Grant P30CA013696 and used the Genomics and High Throughput Screening Shared Resource.

