## Supplementary Figures and Tables for "Deconvolution of Cell Type-Specific Drug Responses in Human Tumor Tissue with Single-Cell RNA-seq"

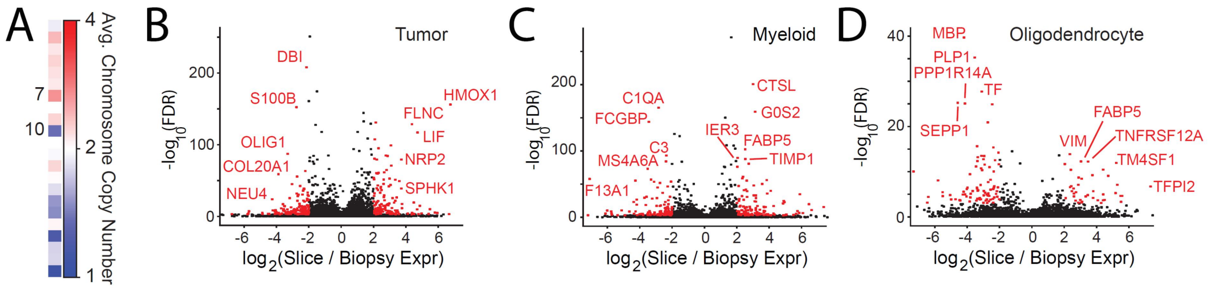


**Figure S1**. A) Heatmap showing the average chromosomal copy number from whole genome sequencing of PW032. B) Volcano plot of differential expression analysis between slice culture and biopsy specimens for the transformed tumor cells in PW032. Genes highlighted in red have FDR<0.05 and absolute fold-change > 4. C) Same as B) for the myeloid cells in PW032. D) Same as B) for the oligodendrocytes.

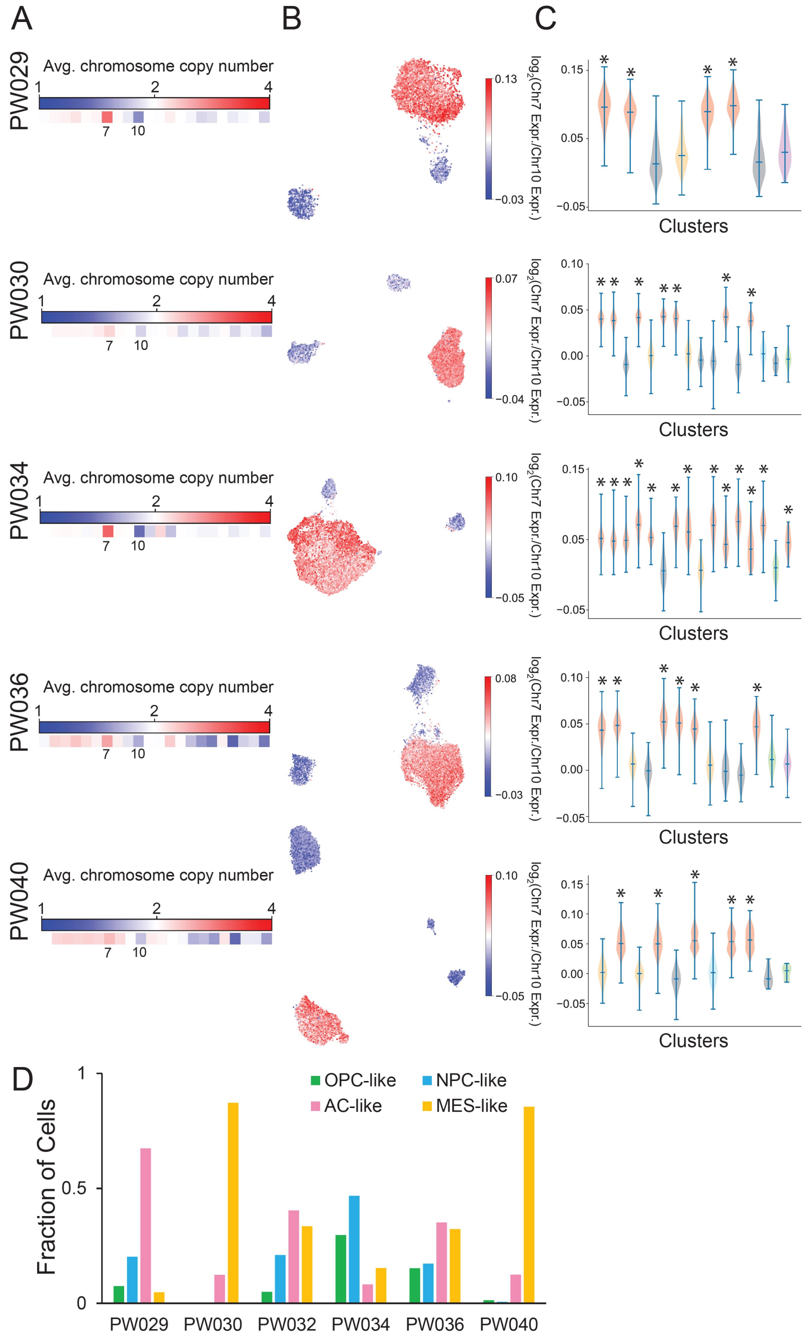


**Figure S2**. A) Heatmaps showing the average chromosomal copy number from whole genome sequencing of PW029, PW030, PW034, PW036, and PW040. B) UMAP embeddings of scRNA-seq profiles from PW029, PW030, PW034, PW036, and PW040 slice cultures colored by the log-ratio of Chr. 7 to Chr. 10 average expression where a high ratio (red) indicates malignant transformation. C) Violin plots showing the distributions of log-ratios of Chr. 7 to Chr. 10 average expression for each Phenograph cluster identified for PW029, PW030, PW034, PW036, and PW040 slice cultures. D) Fractional abundance of each of the four major GBM transformed cell states from the two-dimensional projection in **Fig. 1J** for the transformed cells in the PW029, PW030, PW032, PW034, PW036, and PW040 vehicle-treated slice cultures.

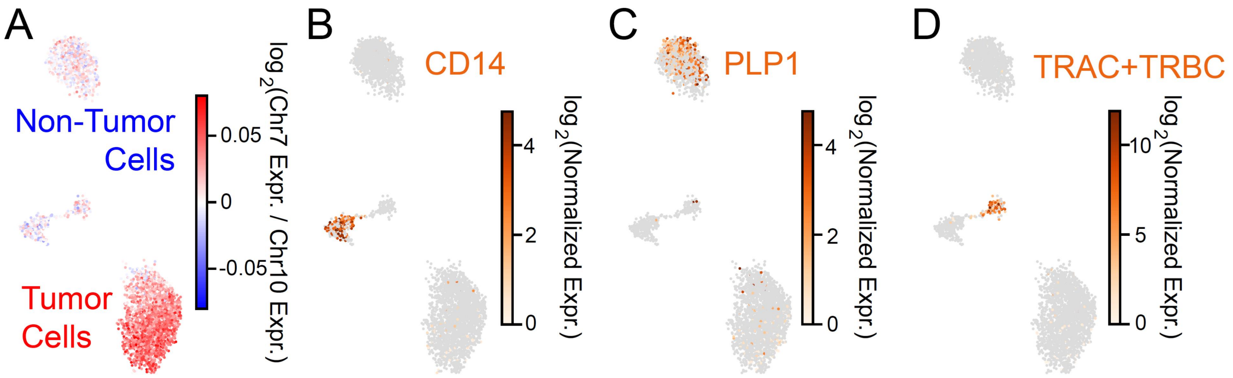


**Figure S3**. A) UMAP embedding of scRNA-seq profiles from five untreated slice cultures taken within 3.5 mm of each other from PW040 colored by the log-ratio of Chr. 7 to Chr. 10 average expression where a high ratio (red) indicates malignant transformation (same as UMAP in **Fig. 1K**). B) Same as A) colored by expression of the myeloid marker CD14. C) Same as A) colored by expression of the oligodendrocyte marker PLP1. D) Same as A) colored by total expression of the T cell receptor constant regions (TRAC, TRBC1, TRBC2).

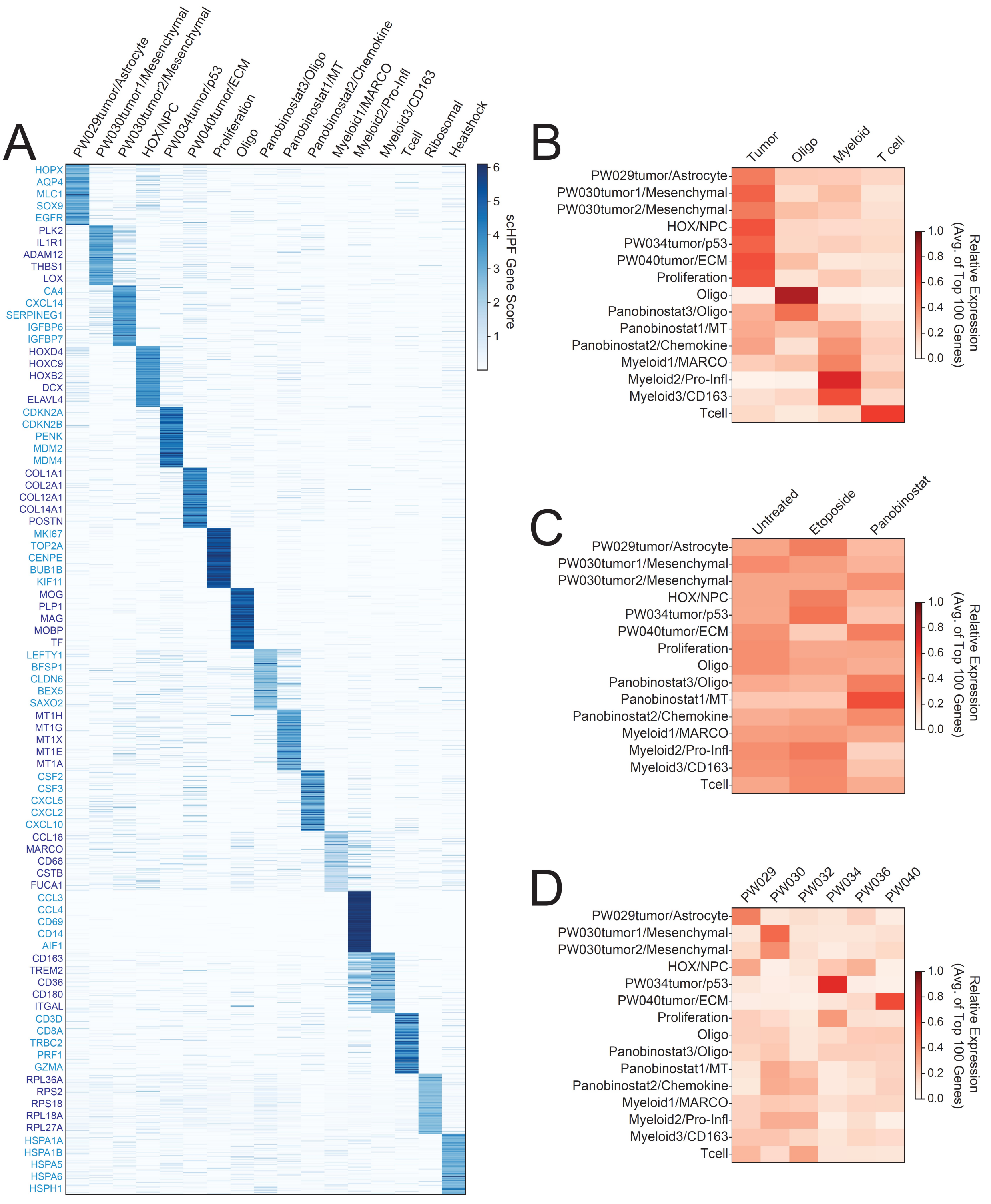


**Figure S4**. A) Heatmap showing the scHPF gene scores for representative high-scoring marker genes for each scHPF factor for the model in **Fig. 3**. B) Heatmap showing the relative average expression of the top 100 markers in each scHPF factor for the transformed (tumor) cells, oligodendrocytes, myeloid cells, and T cells. C) Same as B) for all cells in the vehicle-, etoposide-, and panobinostat-treate slices. D) Same as B) for all cells from all slices for each patient.

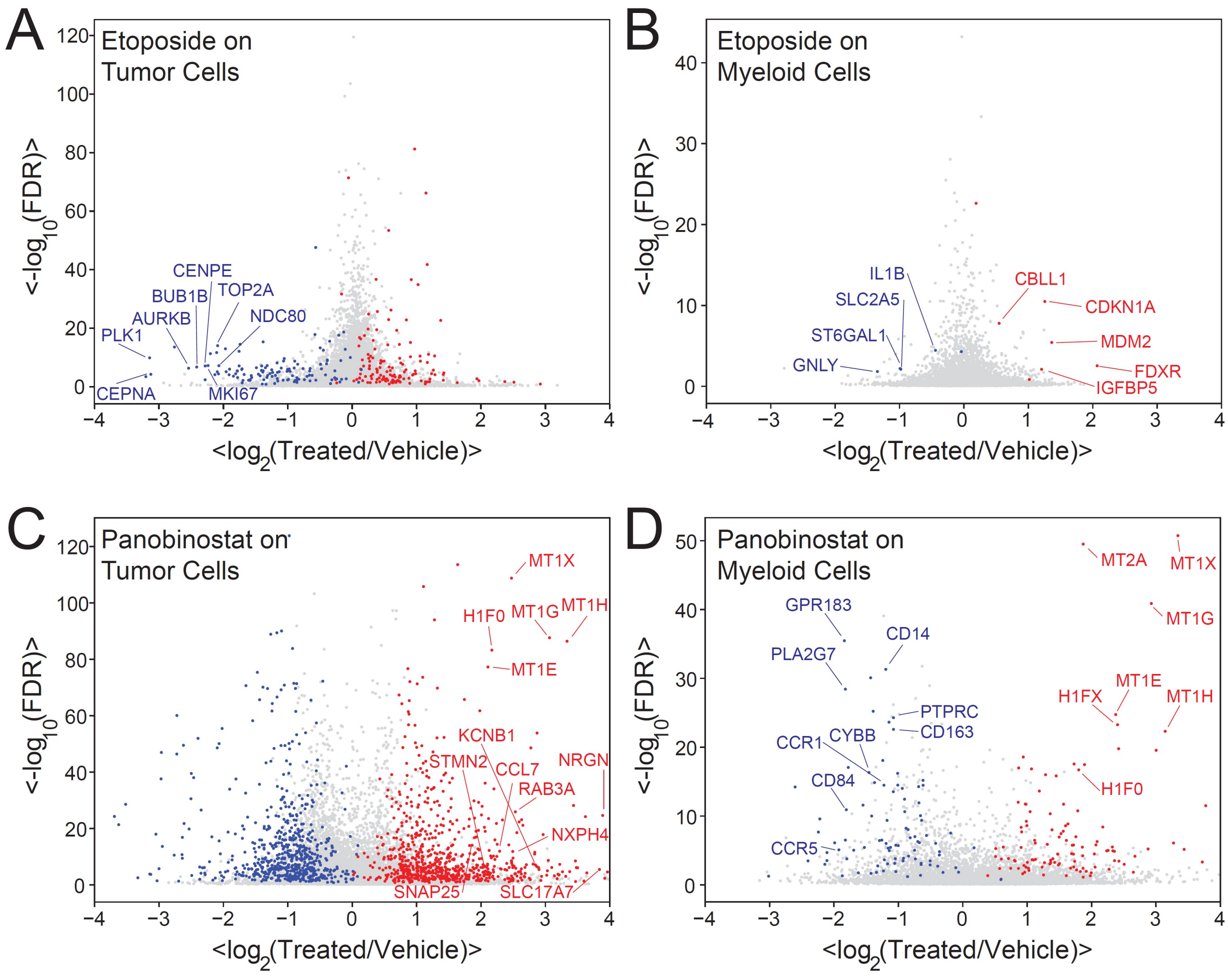


**Figure S5**. A) Volcano plot showing differential expression between etoposide- and vehicle-treated transformed tumor cells average across all patients. Genes highlighted in red and blue have fold-increase or decrease, respectively, greater than two and FDR<0.05 in at least 3/5 patients. A large set of cell cycle control markers are highly downregulated in the etoposide-treated cells. B) Same as A) but for the myeloid cells where etoposide has a smaller effect. C) Same as A) but for panobinostat-treated transformed tumor cells showing strong induction of metallothioneins and several mature neuronal markers. D) Same as C) but for the myeloid cells showing downregulation of the macrophage marker CD163 and strong induction of metallothioneins.

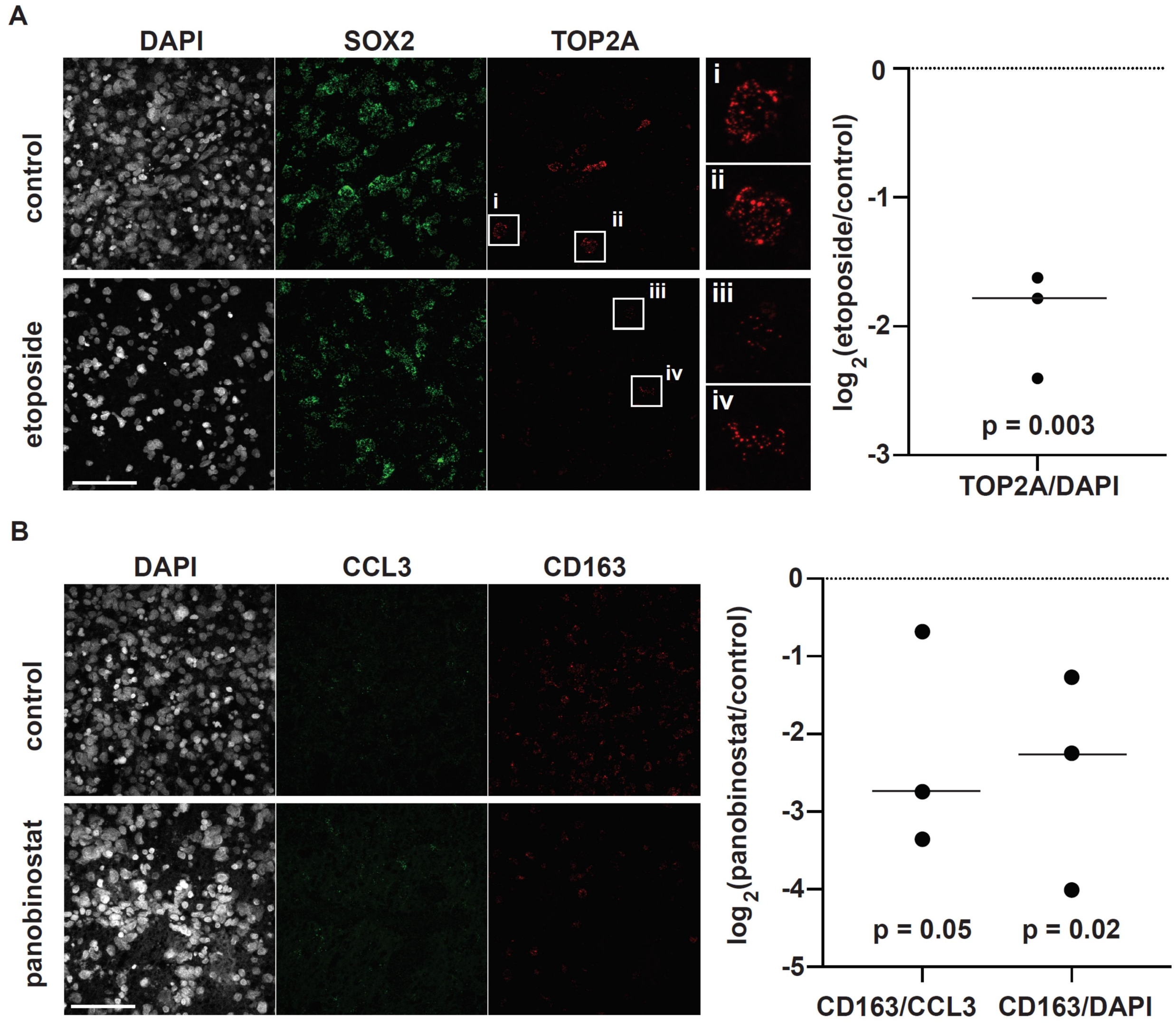


**Figure S6.** A) Representative images (scale bars = 100 microns) of vehicle- and etoposide-treated slices with nuclear stain (DAPI), SOX2 RNAscope, and TOP2A RNAscope demonstrating a loss of TOP2A after etoposide treatment. The plot shows the fold-change in DAPI-normalized TOP2A RNAscope intensity between vehicle- and etoposide-treated slice cultures from three patients (average fold-decrease ~3.7, p = 0.003 from paired t-test). B) Representative images (scale bars = 100 microns) of vehicle- and panobinostat-treated slices with nuclear stain (DAPI), CCL3 RNAscope, and CD163 RNAscope demonstrating a loss of CD163 after panobinostat treatment. The plot shows the fold-change in both DAPI-normalized and CCL3-normalized CD163 RNAscope intensity between vehicle- and panobinostat-treated slice cultures from three patients (average fold-decrease for CD163/CCL3 ~3.4, p = 0.05; average fold-decrease ~4.4 for CD163/DAPI, p = 0.02) demonstrating a consistent loss of CD163 relative to all cells and relative to CCL3.

**Supplementary Tables**

| **Drug Name** | **Resource** | **Cat. #** | **Working Concentration** |
|---|---|---|---|
| Etoposide | Tocris Bioscience | 1226/100 | 2.5 µM |
| Panobinostat(LBH589) | Selleck Chem | S1030 | 0.2 µM |
| Ana-12 | TOCRIS | 4781 - 10 mg | 40 nM |
| Ispinesib | Selleck Chem | S1452 | 1.8 nM |
| Tazemetostat(EPZ-6438) | Selleck Chem | S7128 | 50 µM |
| RO492997 | Selleck Chem | S1575 | 50 nM |

**Table S1.** Summary of drugs used for these studies.

| Sample | Age | Sex | Location | Diagnosis | IDH1 Status | EGFR status | Samples | Fig. 3 Samples |
|---|---|---|---|---|---|---|---|---|
| PW029 | 52 | F | splenial glioma extension into left parietal | Glioblastoma, WHO grade IV | wt | amplified | 1 vehicle slice, 1 etoposide slice | 1 vehicle slice, 1 etoposide slice |
| PW030 | 65 | M | right parietal | Glioblastoma, WHO grade IV | wt | unamplified | 2 vehicle slices, 1 etoposide, 1 panobinostat, 1 ana-12, 1 ispinesib, 1 tazemetostat, and 1 RO492997 slice | 2 vehicle slices, 1 etoposide slice, 1 panobinostat slice |
| PW032 | 61 | M | left frontal | Glioblastoma, WHO grade IV | wt | amplified | 3 uncultured biopsies, 3 vehicle slices, 1 etoposide slice, 1 panobinostat slice | 2 vehicle slices, 1 etoposide slice, 1 panobinostat slice |
| PW034 | 68 | F | left parieto-occipital | Glioblastoma, WHO grade IV | wt | unamplified | 2 vehicle slices, 1 etoposide slice, 1 panobinostat slice | 2 vehicle slices, 1 etoposide slice, 1 panobinostat slice |
| PW036 | 56 | M | right temporal | Glioblastoma, WHO grade IV | wt | amplified | 2 vehicle slices, 1 etoposide slice, 1 panobinostat slice | 2 vehicle slices, 1 etoposide slice, 1 panobinostat slice |
| PW040 | 69 | M | right temporal | Glioblastoma, WHO grade IV | wt | amplified | 5 vehicle slices, 1 panobinostat slice | 2 vehicle slices, 1 panobinostat slice |
| TB6186 | 66 | M | right parietal | Glioblastoma, WHO grade IV | wt | amplified | 1 vehicle slice, 1 etoposide slice, 1 panobinostat slice **(Figure S6)** | |
| TB6193 | 67 | M | left frontal | Glioblastoma, WHO grade IV | wt | unamplified | 1 vehicle slice, 1 etoposide slice, 1 panobinostat slice **(Figure S6)** | |
| TB6199 | 49 | M | right frontoparietal | Glioblastoma, WHO grade IV | wt | amplified | 1 vehicle slice, 1 etoposide slice, 1 panobinostat slice **(Figure S6)** | |

**Table S2.** Summary of patients and specimens used for scRNA-seq.

| Target Gene | Species | Probe | Product Name | Catalog number |
|---|---|---|---|---|
| TOP2A | Homo Sapiens | C1 | RNAscope® Probe- Hs-TOP2A | 470321 |
| SOX2 | Homo Sapiens | C3 | RNAscope® Probe-Hs-SOX2-C3 | 400871-C3 |
| CD163 | Homo Sapiens | C1 | RNAscope® Probe-Hs-CD163 | 417061 |
| CCL3 | Homo Sapiens | C3 | RNAscope® Probe-Hs-CCL3-C3 | 455331-C3 |

**Table S3.** Summary of RNAscope probes used for validation studies.

**Table S4.** Gene score matrix for each scHPF factor in the model shown in **Fig. 3** (attached as a Microsoft Excel sheet).
